# The Microglial Protein sTREM2 Inhibits the Bacterial Functional Amyloid CsgA and Suppresses Amyloid-Dependent Biofilm Formation

**DOI:** 10.64898/2026.07.03.736422

**Authors:** Anthony Balistreri, Mark Gomulinski, Matthew R. Chapman, Jeffery W. Kelly

**Affiliations:** Department of Chemistry, The Scripps Research Institute, La Jolla, CA 92037, USA; Department of Molecular, Cellular and Developmental Biology, University of Michigan, Ann Arbor, MI 48109, USA

**Author notes:** Matthew Chapman and Jeffery Kelly are co-corresponding authors. **Author Contributions:** Conceptualization: AB; Formal Analysis: AB; Funding acquisition: MRC, JWK; Investigation: AB, MG; Methodology: AB; Project administration: AB; Supervision: MRC, JWK; Validation: MRC, JWK; Visualization: AB; Writing – original draft: AB; Writing – review & editing: AB, MG, MRC, JWK.

**Keywords:** CsgA, TREM2, Amyloid Inhibition, Bacterial Biofilm

## Abstract

Protein misfolding and aggregation, including amyloid fibril formation, underlie a large class of human diseases including prominent neurological disorders such as Alzheimer’s and Parkinson’s disease. A small number of human proteins have been identified that inhibit amyloidogenesis. One such protein is sTREM2, a soluble receptor liberated from microglia, the resident macrophages of the central nervous system. The extracellular domain of TREM2 is shed upon proteolytic cleavage to create sTREM2, which has previously been shown to inhibit amyloid-β aggregation *in vitro*. TREM2 is also expressed by intestinal macrophages, which are known to directly bind the bacterial amyloid curli and mount cytokine responses upon exposure. Here we show that sTREM2 is a sub-stoichiometric inhibitor of CsgA amyloidogenesis, CsgA being the major protein component of curli that drives biofilm formation in uropathogenic *Escherichia coli* and many other proteobacteria. *In vitro*, sTREM2 potently and sub-stoichiometrically inhibited CsgA amyloidogenesis in a dose-dependent manner. Kinetic modeling indicated that sTREM2 slowed primary and secondary nucleation, rather than altering fiber elongation. When added exogenously to bacterial growth medium, sTREM2 significantly suppressed curli-dependent pellicle biofilm formation without affecting bacterial growth. These findings establish sTREM2 as a member of the small group of human proteins capable of inhibiting bacterial functional amyloidogenesis, suggesting that gut-resident TREM2-expressing macrophages, which are already known to interact with curli, may employ sTREM2 as a physiologically relevant defense against bacterial amyloid formation.

## Introduction

The misfolding and misassembly of proteins into highly stable amyloid fibril structures is a hallmark of several of the most devastating degenerative diseases, including Alzheimer’s disease (AD) and Parkinson’s disease (PD). Protein misfolding diseases are a large class of hereditary and sporadic disorders that remain poorly understood and only partially addressed by current therapeutics (1).

A small number of proteins have been identified that interact directly with amyloidogenic proteins, inhibiting their aggregation (2–4). One such protein is TREM2, triggering receptor <u>e</u>xpressed on <u>m</u>yeloid cells <u>2</u>, a receptor protein most notably expressed by microglia, the resident macrophages of the central nervous system (5, 6). TREM2 regularly undergoes cleavage to shed the receptor’s ectodomain, termed soluble TREM2 or sTREM2, into the extracellular space (7, 8). In 2021, Vilalta et al. was first to describe sTREM2 as an effective inhibitor of amyloid-β aggregation *in vitro* (9).

TREM2 is not expressed exclusively in the brain. TREM2-expressing macrophages are also present in the intestine (10, 11), where they are known to directly interact with bacterial amyloid fibrils called curli (12). Curli is the major amyloid component of biofilms formed by Escherichia coli and other enteric bacteria (13), and is recognized as a pathogen-associated molecular pattern by the intestinal immune system (12). This raised the question of whether sTREM2, in addition to its established activity against amyloid-β, might also act directly on curli or CsgA, the secreted, amyloid-forming protein that is the major structural subunit of curli fibers.

Proteins that display anti-amyloid activity have common immunoglobulin folds like the v-set or β-sandwich fold. Two such proteins that share the β-sandwich fold are the bacterial amyloid inhibitor protein CsgC (14) and the human retinol transport protein transthyretin (TTR) (15) (**Figure 1A**). CsgC is a small chaperone-like protein and member of an *E. coli* operon that controls the formation of curli (16). CsgC inhibits amyloid formation of its native client CsgA (17). Though TTR is known to form amyloid itself (18), stabilization of the monomeric (M-TTR) form of the protein allows the monomer to act as an amyloid inhibitor (19). Just like CsgC, M-TTR is also an effective inhibitor of CsgA aggregation (19). When we examined the structure of sTREM2 and found it to be similar to CsgC and M-TTR, we asked whether sTREM2 shared the same amyloid inhibitor activity.

**Figure 1.**
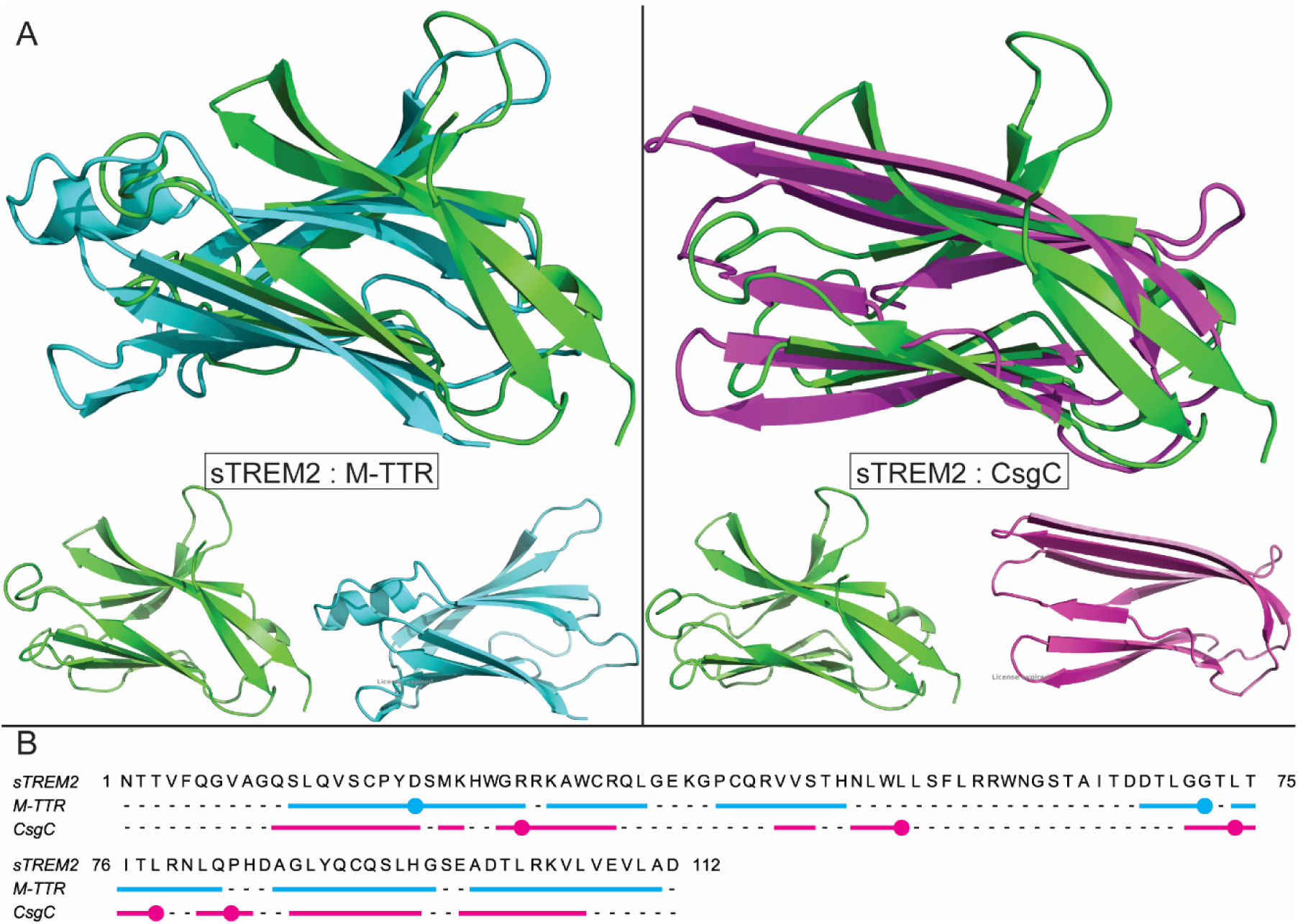
Structural comparison between sTREM2, M-TTR, and CsgC. **A)** sTREM2 (PDB:5ELI, green) was superimposed with M-TTR (PDB: 1GKO, cyan) and CsgC (PDB:2Y2Y, magenta) using the SSM algorithm. A summary of the results and additional alignment measurements can be found in **Table 1. B)** A structural alignment showing the sTREM2 amino acid sequence. The colored bars correspond to the residues in sTREM2 that superimpose with M-TTR (cyan) and CsgC (magenta). Circles describe wherever the aligned amino acids are identical between proteins (M-TTR = 2/112; CsgC = 5/112). Full sequences of each protein and the assigned secondary structure to each amino acid can be found in **Supplementary Figure 1**.

Here we show that sTREM2 is a sub-stoichiometric inhibitor of CsgA aggregation. When sTREM2 is added to purified CsgA, there is a dose-dependent decrease in ThT binding and fibril formation. Kinetic modeling indicates that sTREM2 inhibits CsgA aggregation by slowing primary and secondary nucleation. When added exogenously to bacterial culture medium, sTREM2 is an effective inhibitor of CsgA-dependent biofilm formation.

## Results

### sTREM2 is an amyloid inhibitor protein

sTREM2 shares a common structural motif with other putative amyloid inhibitor proteins like CsgC and M-TTR (**Figure 1A**, **Table 1**). The proteins CsgC and M-TTR are both capable of efficient inhibition of amyloid formation from the bacterial functional amyloid CsgA (16, 19, 20). sTREM2, CsgC, and M-TTR share a β-sandwich V-type Immunoglobulin-like fold (**Figure 1A**, **Table 1)**. Structural similarity between sTREM2:M-TTR and sTREM2:CsgC was assessed using three independent computational methods (**Table 1**). PDBeFold identified marginal but statistically non-random alignments for both pairs (sTREM2:M-TTR: Q = 0.137, Z = 2.178, RMSD = 3.99 Å; sTREM2:CsgC: Q = 0.134, Z = 2.299, RMSD = 3.29 Å). TM-align, which uses a length-normalized scoring function less sensitive to alignment size, yielded TM-scores of 0.571 and 0.534 for the M-TTR and CsgC comparisons respectively, both exceeding the established same-fold threshold of 0.5 (21). The same-fold assignment was further supported by GDT_TS scores of 47.8% and 45.8% and MaxSub scores of 0.705 and 0.663, indicating that approximately 70% of residues in each comparison superpose within a structurally meaningful range, despite the low sequence identity across aligned residues (2–5%) (**Figure 1B, Figure S1**).

**Table 1.**
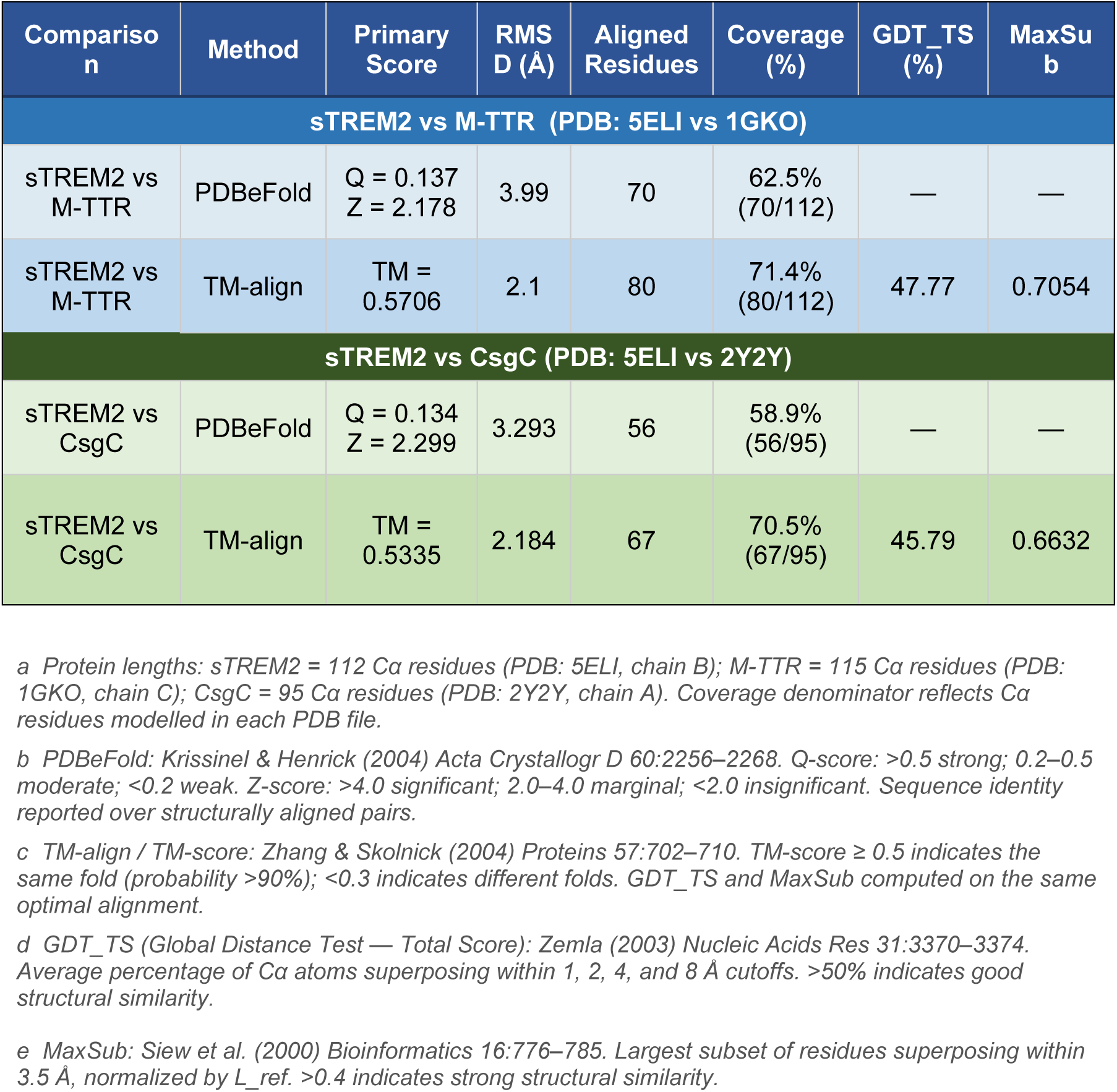
Structural alignment of sTREM2 with M-TTR and CsgC quantified by multiple independent methods.

sTREM2 is a sub-stoichiometric inhibitor of CsgA amyloid fibril formation in a ThT fluorescence assay. When mixed with ThT in phosphate buffer, CsgA spontaneously formed ThT-positive aggregates within 4 hours (Figure 2A) having a fibril morphology (*vide infra*) (22). When purified sTREM2 was added to freshly purified CsgA protein, sTREM2 increased the ThT lag phase of CsgA in a dose-dependent manner (**Figure 2A-B**). Indeed, sTREM2 fully inhibited aggregation in the time course of the assay at a 1:10 sTREM2:CsgA stoichiometric ratio (**Figure 2A**).

**Figure 2.**
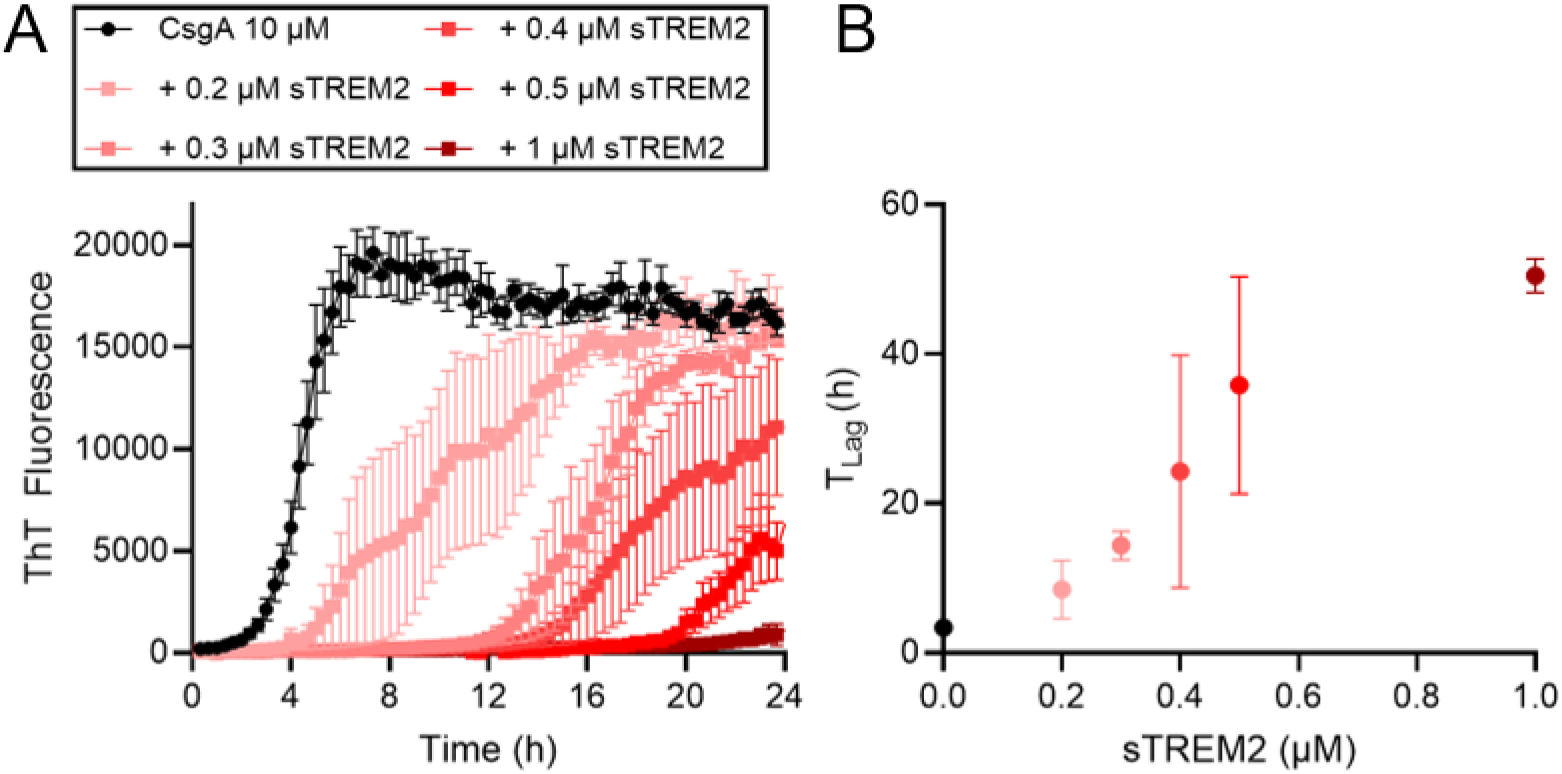
sTREM2 inhibits CsgA amyloid formation in a dose-dependent manner. **A)** ThT fluorescence aggregation assays of freshly purified CsgA (10 µM) in the presence of increasing concentrations of sTREM2 (0 to 1 µM). Curves represent the mean of triplicate experiments and error bars show SEM. The progressive rightward shift and reduction in final fluorescence signal with increasing sTREM2 concentration reflect dose-dependent inhibition of both the kinetics and extent of CsgA amyloid formation. **B)** Quantification of the lag phase (T_Lag_) for each condition was determined by fitting individual replicate ThT curves to a logistic sigmoidal function as described by Arosio et al (36). Data points represent the mean of triplicate experiments and error bars show SEM.

### sTREM2 inhibits primary and secondary nucleation to inhibit amyloidogenesis

Amyloid fiber formation is a complex, multi-step process that relies on several steps and each step can be modeled through rate constants of formation (23). Amyloid fibril formation proceeds through primary nucleation (k_n_), elongation (k_+_), and under some conditions secondary nucleation (k_2_), and Amylofit software can identify the step(s) most affected by an inhibitor through global fitting of ThT kinetic data (23). To determine which step sTREM2 inhibits, ThT aggregation assays were performed at 0, 2%, and 10% seed concentrations with varying sTREM2 concentrations at each condition (**Figure 3**). Because high-seed reactions (10% w/w, **Figure 3E-F**) are dominated by elongation (k_+_) while low-seed (2% w/w, **Figure 3C-D**) and unseeded reactions (**Figure 3A-B**) reflect primary and secondary nucleation (k_n_ and k_2_), this design allows Amylofit to globally fit the data against mechanistic models of inhibition and identify the best-fitting model by lowest mean residual error (23). When we performed the ThT assays across a seed concentration series described above, the model consistent with inhibition of primary and secondary nucleation provided the best fit across conditions for CsgA (**Figure 3B, D, and F**).

**Figure 3.**
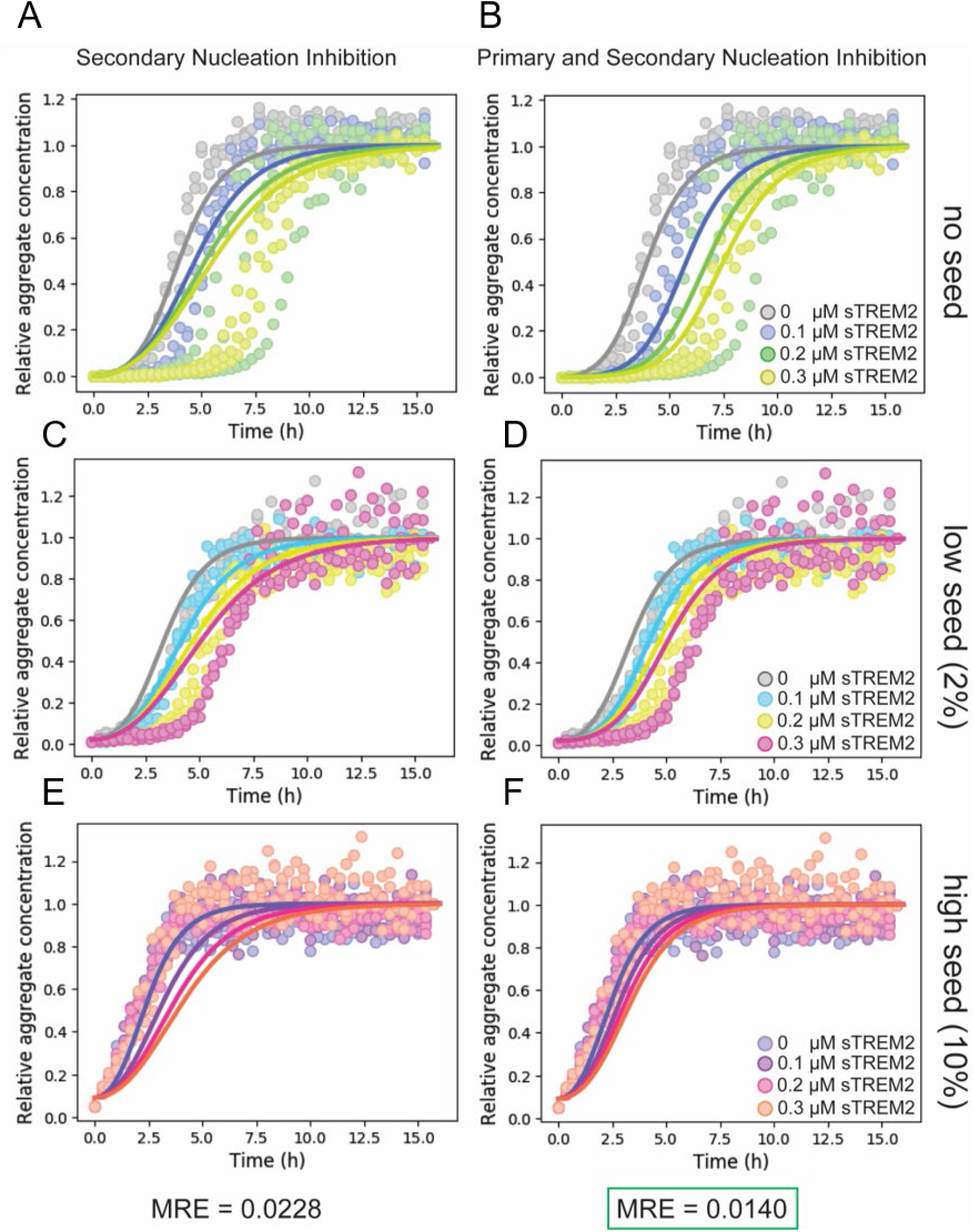
Amylofit kinetic modeling identifies primary and secondary nucleation as the mechanism of sTREM2-mediated inhibition of CsgA aggregation. **A-F)** ThT fluorescence aggregation assays of CsgA monomer (10 µM) with increasing concentrations of sTREM2 (0, 0.1, 0.2, and 0.3 µM) performed under three seeding conditions: no seed (**A, B**), low seed (2% w/w preformed CsgA fibrils; **C, D**), and high seed (10% w/w preformed CsgA fibrils; **E, F**). All assays were performed in triplicate; data points represent individual replicate measurements. Each curve is independently normalized to its own minimum and maximum fluorescence values. Solid lines represent fits to a secondary nucleation-dominated aggregation model using Amylofit software. The left column (**A, C, E**) shows fits to a model in which sTREM2 inhibits secondary nucleation only; the right column (**B, D, F**) fits to a model in which sTREM2 inhibits both primary and secondary nucleation. The mean residual error (MRE) for each model is shown below the respective column. The lower MRE for the primary and secondary nucleation model (MRE = 0.0140, green box) compared to the secondary nucleation-only model (MRE = 0.0228) indicates that inhibition of both primary and secondary nucleation provides the best description of sTREM2’s mechanism of action.

### The addition of sTREM2 to CsgA leads to decreased fibril formation

In the presence of CsgC or M-TTR, CsgA shows diminished propensity to form amyloid fibers visible by TEM (16, 19). A comparable reduction in fiber formation was observed when CsgA was challenged with sTREM2 (**Figure 4**). In the absence of any inhibitor, CsgA readily formed amyloid fibers visible in large, uniform bundles in TEM micrographs (**Figure 4A**). sTREM2 was added to CsgA at the same ratios tested in the ThT assays shown in **Figure 2**. As the amount of sTREM2 added to the amyloidogenic protein increased, the amount of amyloid fibers observed on the TEM grid decreased (**Figure 4B-C**), consistent with the decrease in observed ThT fluorescence signal (**Figure 2**).

**Figure 4.**
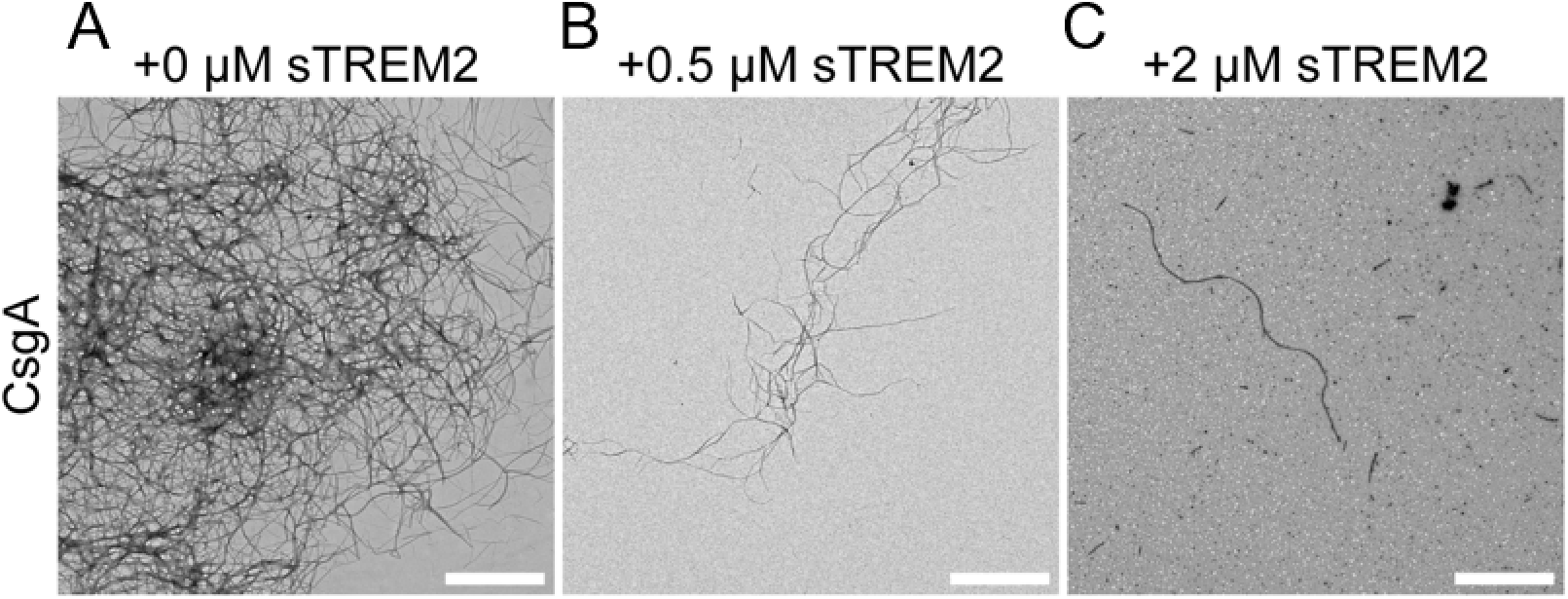
CsgA fibril formation is decreased in the presence of inhibitory levels of sTREM2. Samples were taken during CsgA ThT binding assays. sTREM2 was added to freshly purified CsgA at **A-C)** 0 µM, 0.5 µM, and 2 µM. Samples were taken at 8 hours, spotted on TEM grids, and the grids were stained with uranyl acetate and visualized by TEM. The scale bars represent 10 µm.

### sTREM2 inhibits amyloid-dependent biofilm formation

Bacterial biofilms are collections of bacteria cells enmeshed in an extracellular matrix composed of polysaccharides, nucleic acids, and bacterially-derived amyloid fibrils (24). In the case of uropathogenic *E. coli*, the pellicle biofilm it produces at the air-liquid surface requires the secretion and aggregation of CsgA into amyloid fibrils (25). Given the amyloid inhibition activity of sTREM2 *in vitro* (*vide supra*), we wanted to explore the exogenous addition of sTREM2 to a bacterial culture under biofilm formation conditions to see if we could achieve biofilm inhibition. UTI89 is a well-characterized uropathogenic strain of *E. coli* that robustly forms curli-dependent pellicle biofilms under standard biofilm-inducing conditions (25). We supplemented the culture medium of *E. coli* UTI89 with 0.1, 0.5, and 2 µM sTREM2 and after 48 hours checked for biofilm formation (**Figure 5**). Images of the biofilm taken from above showed a decrease in the wrinkled pellicle biofilm when sTREM2 was added to the culture medium, and no visible pellicle biofilm was seen in wells that contained 2µM of sTREM2 (**Figure 5A**). As a control, a UTI89 strain that lacks the *csgA* gene was also unable to form visible pellicle biofilms (**Figure 5A**). In the same concentrations tested in the biofilm assay, sTREM2 did not affect cell growth of either bacterial strain (**Figure S2**). Crystal violet dye stains biofilm mass and a measurement of crystal violet absorbance (A595) can provide a semi-quantitative assay for measuring biofilm (25). Using the crystal violet assay, sTREM2 supplementation significantly decreased biofilm mass, confirming a decrease in amyloid-dependent biofilm formation (**Figure 5B**). Before staining, small samples of bacteria from a few conditions were imaged by TEM which showed a decrease in visible fibers associated with the cells (**Figure 5C**). These data suggest that sTREM2 is capable of inhibiting amyloid formation in a disease-relevant *in vivo* environment.

**Figure 5.**
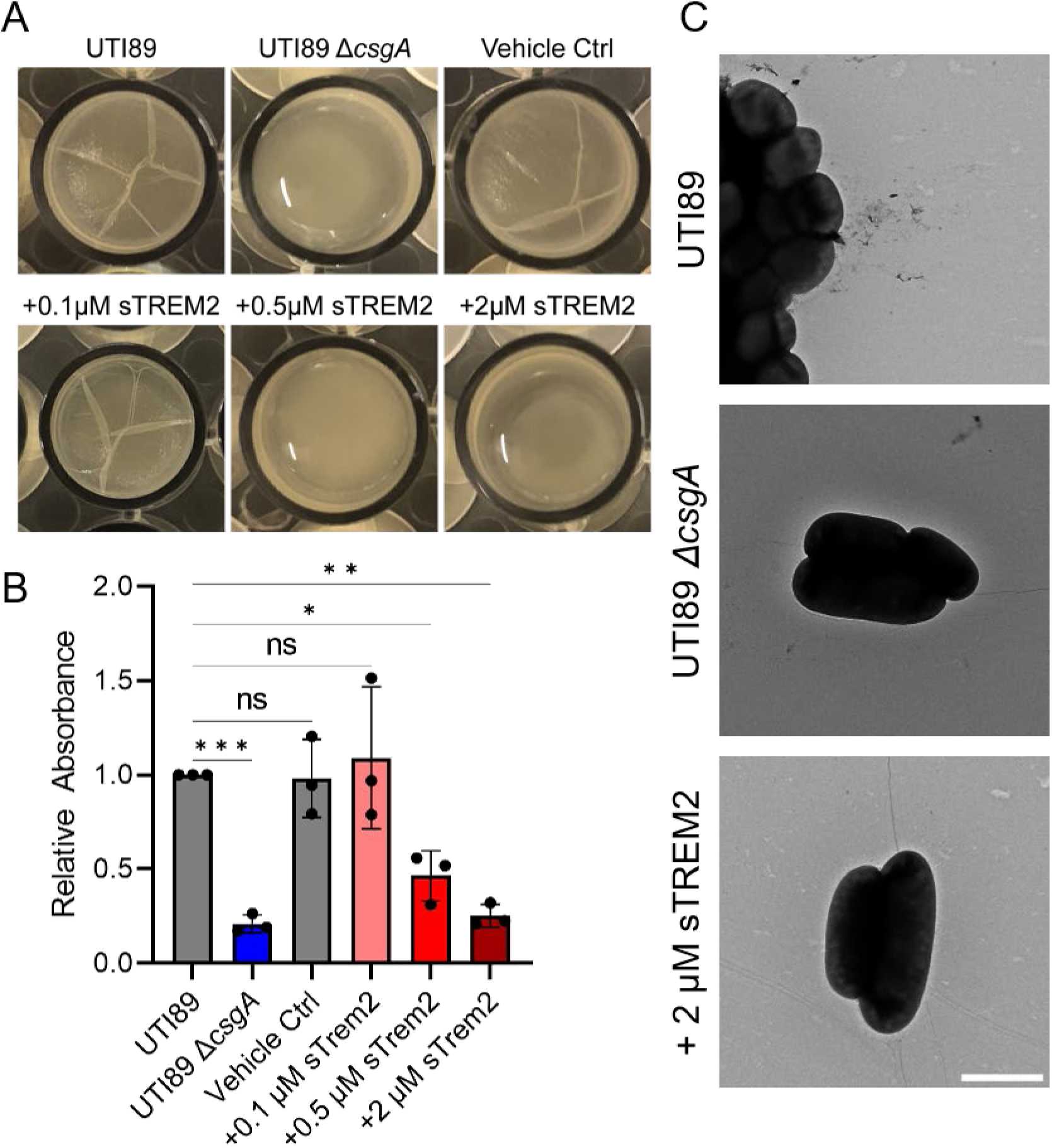
sTREM2 can inhibit amyloid-dependent biofilm formation. The uropathogenic *E. coli* strain UTI89 and a ΔcsgA knockout were cultured under biofilm inducing conditions. **A)** After 48 hours pictures were taken of each well that had sTREM2 supplemented (or a vehicle only control) medium at the labeled concentration. **B)** The pellicle biofilm was stained with crystal violet dye and dissolved with concentrated acetic acid. The A595 from each condition is shown relative to the bacteria only negative control. Data are presented as mean ± SEM (n = 3 per condition); differences among conditions were assessed by one-way ANOVA (F(5, 12) = 14.02, P = 0.0001) with Dunnett’s post hoc multiple comparisons test. Significant reductions relative to UTI89 were observed for the ΔcsgA control (***P = 0.0010), 0.5 μM sTREM2 (*P = 0.0167), and 2 μM sTREM2 (**P = 0.0015); the vehicle control and 0.1 μM sTREM2 did not differ significantly. **C)** Samples from the labeled conditions were spotted onto carbon/formvar TEM grids and negatively stained to visualize bacteria and their associate fibers. Representative micrographs are shown and scale bar represents 10 µm.

## Discussion

### Multi-substrate amyloid inhibition

sTREM2 is an inhibitor of functional bacterial amyloid fibril formation from the CsgA protein. sTREM2 can be added to a growing list of proteins capable of inhibiting amyloid formation across multiple substrates, including CsgC (16), M-TTR (19), Gkn1 (26), *H. pylori* virulence factor CagA (27). Many amyloid inhibitor proteins share a 3D structure, the beta-sandwich or V-type immunoglobulin-like fold (**Figure 1**). The immunoglobulin-like fold is common; the largest group of human proteins by folding domain is the immunoglobulin-like fold (28, 29). The V-type immunoglobulin-like domain takes its name from the antibody variable domain, the structural module responsible for epitope recognition in adaptive immunity (30). Whether sTREM2 binds amyloid proteins through an analogous specific interaction or through a more general recognition of the cross-β-sheet architecture common to all amyloid fibers remains to be determined. The two amyloids sTREM2 has been shown to inhibit *in vitro* — CsgA and amyloid-β — share little sequence similarity but they both form fibers with a cross-β-sheet architecture, suggesting the latter possibility is more likely.

sTREM2 inhibited amyloid formation by slowing primary and secondary nucleation (**Figure 3**). Nucleation inhibition is increasingly recognized as a common mechanism among amyloidogenesis inhibitor proteins: Nagaraj et al. showed that molecular chaperones suppress primary nucleation during functional amyloid formation (2), and inhibition of nucleation has also been reported for other small amyloid inhibitor proteins including CsgC (20). For sTREM2 specifically, this mechanism has precedent in the Aβ literature: Belsare et al. demonstrated that sTREM2 binds fibrillar Aβ and selectively inhibits secondary nucleation (31). Our findings extend this picture to an additional amyloid substrate, suggesting that nucleation inhibition may be a general feature of sTREM2’s multi-substrate inhibitory activity rather than a substrate-specific effect.

### Biofilm inhibition

Curli-dependent biofilms are associated with persistent uropathogenic infections and represent a significant clinical challenge, as biofilm-encased bacteria are substantially resistant to antibiotics and immune clearance (32). Biofilms contribute to infectious disease by allowing bacterial colonies to grow under environmental hazards that would normally be toxic to them (33). Many bacterial biofilms rely on the secretion and aggregation of functional amyloid proteins that self-assemble into amyloid fibrils (34). When sTREM2 was exogenously added to UTI89 growth medium, the effect was a decrease in biofilm production without an apparent effect on the growth of the bacteria (**Figure 5, Figure S2**). That an endogenous human immune protein can suppress curli-dependent biofilm formation without affecting bacterial growth extends a precedent established by TTR (19), and raises the possibility that sTREM2 or structurally similar immunoglobulin-fold proteins could inform a new class of anti-biofilm agents targeting functional amyloid.

We have demonstrated that sTREM2 is a sub-stoichiometric inhibitor of CsgA, acting through suppression of primary and secondary nucleation, and that it suppresses curli-dependent biofilm formation in a uropathogenic *E. coli* model. Together with the prior demonstration that sTREM2 inhibits amyloid-β aggregation (9), these findings suggest that sTREM2 could have multi-substrate anti-amyloid activity across diverse amyloidogenic substrates. These results also have potential implications for Parkinson’s disease pathobiology. Prior work from our group showed that curli-producing bacteria in the gut promote α-synuclein aggregation in the enteric nervous system, accelerating motor deficits in a mouse model of Parkinson’s disease (35). If gut-resident TREM2-expressing macrophages encounter curli *in vivo*, the amyloid inhibitory sTREM2 activity demonstrated here could represent a previously unappreciated mechanism by which the gut innate immune system limits the propagation of amyloid-seeding events that may initiate or accelerate synucleinopathies. Exploring sTREM2 as an inhibitor of gut-to-brain amyloid seeding represents an important and tractable future direction.

## Materials and Methods

Detailed protocols for all methods described below are provided in SI Appendix, Materials and Methods.

### Bacterial Growth and Strain/Cell line List

Overnight bacterial cultures were grown in LB with ampicillin (100 μg/mL) or kanamycin (50 μg/mL) at 37 °C with shaking; strains and cell lines used in this study are listed in SI Appendix, Materials and Methods.

### Protein Purification

CsgA was expressed and purified as described previously. CsgA curli were generated by extended incubation of monomeric protein, and lyophilized human sTREM2 was purchased and stored at -80 °C until use.

### Secondary Structure Analysis

Structures of sTREM2 (PDB: 5ELI), CsgC (PDB: 2Y2Y), and M-TTR (PDB: 1GKO) were superimposed by secondary-structure matching, and the statistical quality of each alignment was assessed using PDBeFold.

### Thioflavin T Binding Assay

Amyloid formation by CsgA was monitored in triplicate in 96-well plates by the increase in thioflavin-T fluorescence (450 nm excitation, 495 nm emission), in the presence or absence of sTREM2 and sonicated seeds, as described previously.

### Transmission Electron Microscopy

Purified protein or bacterial culture samples were spotted onto formvar/carbon-coated copper grids, negatively stained with 2% uranyl acetate, and imaged on a JEOL JEM-1400plus transmission electron microscope, as described previously.

### Amyloid Dependent Pellicle Biofilm Formation

Pellicle biofilms were grown from UTI89 cultures in YESCA medium in 48-well plates, with purified sTREM2 added at the indicated concentrations, and incubated at 26 °C for 2 days before imaging to assess biofilm morphology, as described previously.

### Crystal Violet Assay

Biofilm mass was quantified by staining washed biofilms with 0.1% crystal violet, solubilizing the retained dye in 33% acetic acid, and measuring absorbance at 595 nm, as described previously.

### Quantification and Statistical Analysis

Graphs were produced in GraphPad Prism v10 and data shown is the average of *n* number of technical replicates (as noted in figure legend) with error bars depicting the standard error of the mean. Statistical significance was assigned based on a one-way ANOVA with Dunnett’s post hoc multiple comparisons test using Prism’s default parameters where * denotes p < 0.05, ** denotes p < 0.01, *** denotes p < 0.001, and **** denotes p < 0.0001.

## Acknowledgments

The authors would like to acknowledge all the helpful conversations and contributions of the extended members of the Chapman and Kelly labs, especially Dr. Russ Gibadullin and Divya Kolli. We would also like to acknowledge Georg Miesl for his assistance in correctly utilizing the Amylofit software. This work was supported by NIH grant R01GM118651-09 to MRC and a Freedom Together Foundation grant to JWK.

## Notes

### Competing Interest Statement

The authors have declared no competing interest.

