## Supplemental Figures and Methods for "The Microglial Protein sTREM2 Inhibits the Bacterial Functional Amyloid CsgA and Suppresses Amyloid-Dependent Biofilm Formation"

**This PDF file includes:**

SI Materials and Methods
Figures S1 to S2
Tables S1
SI References

### SI Materials and Methods

#### Bacterial Strains and Growth Conditions

All overnight cultures were grown in LB broth supplemented with 100 μg/mL ampicillin. When required, LB agar plates were supplemented with the same antibiotics at the same concentrations. Strains used in this study are listed in Table S1 below.

**Table S1.** Bacterial and cell strains used in this study.

| Strain | Relevant Notes | References |
|---|---|---|
| CsgA expression strain | NEB 3016 *ΔslyD* + pCsgA (pET11d-CsgA-sec 6xHis) | (1) |
| UTI89 | Wild-type clinical UPEC | (2) |
| *UTI89 ΔcsgA* | Deletion of the major curlin gene csgA in UTI89; abolishes curli expression | (3) |

**Protein Purification.** CsgA was purified as described previously (1). Curli amyloid fibers composed of CsgA were produced by incubating an aliquot of monomeric protein at room temperature for at least 3 days. Lyophilized human sTREM2 (His19-Ser174) with a C-terminal His tag (Cat. No. TR2-H52H5-100ug) was purchased and resolubilized following the provided Certificate of Analysis with deionized water and stored at -80 °C, thawing just before use.

**Secondary Structure Analysis.** The structures of sTREM2 (PDB: 5ELI, chain B), monomeric transthyretin (M-TTR; PDB: 1GKO, chain C), and CsgC (PDB: 2Y2Y, chain A) were retrieved from the RCSB Protein Data Bank. Pairwise structural alignments of sTREM2 with M-TTR and sTREM2 with CsgC were performed using two independent computational methods.
Initial alignments were carried out using PDBeFold (4), which performs secondary-structure-based superposition and reports a Q-score (a composite measure of alignment quality normalized for the number of aligned residues and their root mean square deviation; RMSD), a Z-score reflecting statistical significance relative to random structural comparisons, and the percentage of sequence identity over aligned residue pairs. Structural alignments were independently assessed using the TM-align algorithm (5). Global Distance Test Total Score (GDT_TS; (6)) and MaxSub score (7) were computed on the same optimal TM-align superposition as additional independent measures of structural similarity. GDT_TS reports the average percentage of Cα atoms superposing within distance cutoffs of 1, 2, 4, and 8 Å; MaxSub reports the largest subset of residues superposing within 3.5 Å, normalised by the length of the shorter protein. Structural figures were prepared using PyMOL (v2.5.5, RRID: SCR_000305).

**Thioflavin T Binding Assay.** CsgA assays were performed as previously described (1). Briefly, freshly purified CsgA was diluted with phosphate buffer and combined with an excess of the amyloid-specific dye thioflavin-T (ThT) (Cat No. AC211760250). Amyloid formation was monitored by measuring an increase in ThT fluorescence at 495 nm (450 nm excitation). Assays were performed in triplicate at microscale within 96-well plates and measured with Infinite Nano+ F200 Tecan plate reader. Selected samples include: sTREM2 diluted from a stock solution, CsgA seeds were purified previously and sonicated directly before addition.

**Amylofit Kinetic Modeling.** ThT aggregation assays were performed at a constant CsgA monomer concentration (10 μM) across three seeding conditions (unseeded, 2% w/w, and 10% w/w preformed CsgA fibrils) with varying sTREM2 concentrations (0, 0.1, 0.2, and 0.3 μM) at each seed level. Data were analyzed using Amylofit (8) to globally fit aggregation models incorporating primary nucleation, elongation, and secondary nucleation. The aggregation model used was secondary nucleation dominated. The inhibition model best describing the data was identified by lowest mean residual error (MRE).

**Transmission Electron Microscopy.** TEM images were produced as previously described (9). Briefly, 5 µL of purified protein samples or bacterial culture samples were spotted onto a formvar/carbon 200 mesh copper grid (Cat No.50-260-38). After a 5 min incubation, grids were spotted with 5 µL of DI water followed by 5 µL of 2% uranyl acetate to provide micrograph image contrast. Grids were imaged using a Jeol JEM 1400plus Transmission Electron Microscope.

**Amyloid Dependent Pellicle Biofilm Formation.** Pellicle biofilm assays were performed as previously described (10). Briefly, in wells of a sterile 48-well plate, liquid cultures of UTI89 were started by inoculating 1 mL of sterilized YESCA medium with 1 µL of an overnight culture. Samples of purified sTREM2 were added to the indicated final concentration by diluting from stock solutions. The cultures were incubated at 26 ºC for 2 days. Pictures were taken after the incubation period to visualize pellicle biofilm morphology.

**Crystal Violet Assay**. Semi-quantitative crystal violet assays were performed as briefly described (1). Briefly, after the 2-day incubation period, planktonic bacteria were removed from beneath the biofilm by aspiration. 1 mL of a 0.1% crystal violet solution prepared from a powder stock (Cat. No. V5265) were added to each well and the biofilms incubated in the staining solution for 5 mins. The staining solution was aspirated and the biofilm washed with equal volume of Milli-Q water 3 times. Stained biofilms were dried and dissolved by the addition of an equal volume of 33% acetic acid (glacial acetic acid, diluted to 33% v/v, Cat. No. 695092). A sample from each condition was placed in a Infinite M200 Tecan plate reader and A595 was used to measure biofilm mass.

**Bacterial Growth Curve Assay**. UTI89 and UTI89 ΔcsgA were grown overnight in YESCA medium at 37 °C with shaking. Overnight cultures were diluted into fresh YESCA medium supplemented with sTREM2 at the indicated concentrations or vehicle alone, and optical density at 600 nm (OD600) was monitored over 24 hours in a plate reader with continuous shaking at 37 °C.

### SI Figures

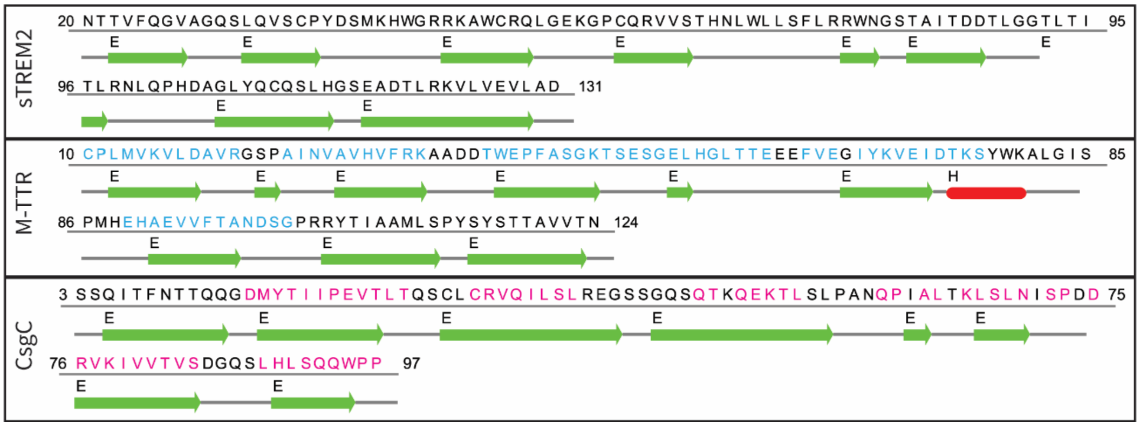


**Figure S1**. **Sequences and secondary structure assignments of TREM2, M-TTR, and CsgC.** The sequence of each protein was extracted from their respective coordinate files, namely: sTREM2, PDB:5ELI; M-TTR, PDB: 1GKO; and CsgC, PDB:2Y2Y. Underneath each sequence is the secondary structure assignment of each amino acid, “E” and a green arrow = Extended β-strand, “H” and a red oval = α-helix. Colored letters are amino acids in each protein that superimpose with sTREM2**.**

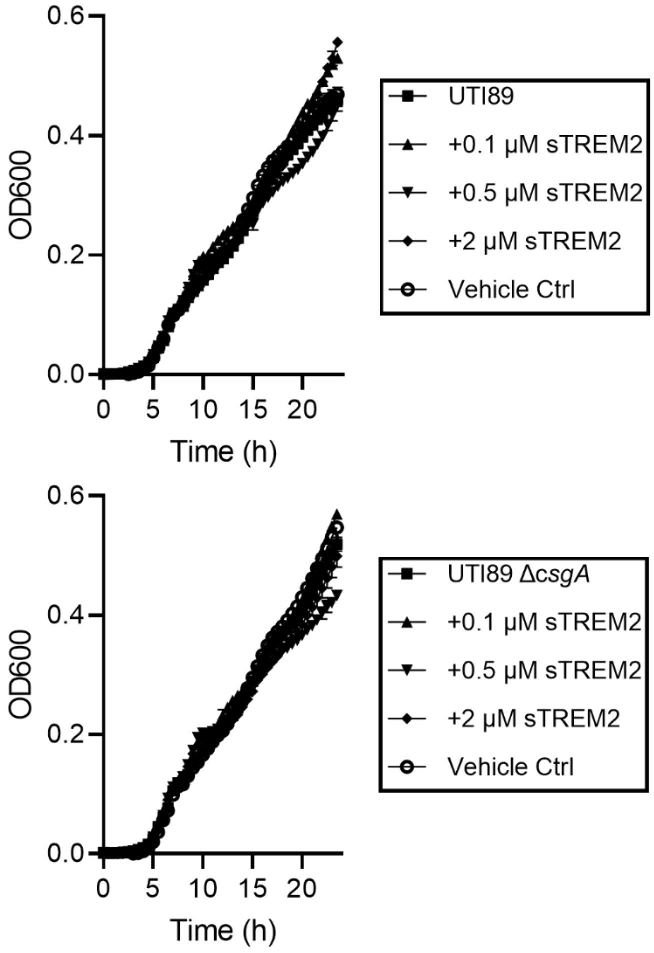


**Figure S2**. **The addition of sTREM2 does not affect cell growth of the biofilm model UTI89.** **A)** UTI89 and **B)** UTI89 ΔcsgA were grown at 37 °C with shaking in YESCA medium supplemented with sTREM2 or vehicle control at the labeled concentrations. The OD600 was monitored in a plate reader for 24 hours and sTREM2 showed no effect on bacterial growth.
